# *Ziziphus spina-christi* as a Refuge and Resource for Parasitoids in Arid Ecosystems

**DOI:** 10.64898/2026.11.12.766100

**Authors:** Norah Saabna, Tamar Keasar

**Affiliations:** Department of Evolutionary and Environmental Biology, University of Haifa, Haifa 3103301, Israel.; Department of Biology and the Environment, University of Haifa - Oranim, Tivon 36006, Israel

**Keywords:** aridity, floral resources, sugar feeding, parasitoid abundance, parasitoid community composition, seasonal variation, refuge habitat, desert afforestation, dryland restoration

## Abstract

1. Climate change is increasing aridity in dryland ecosystems, potentially threatening parasitoid wasps through reduced resource availability and environmental stress. Native drought-adapted plants may help sustain parasitoid communities by providing floral resources and refuge habitats. The tree *Ziziphus spina-christi* (L.) tolerates drought and heat stress, making it a promising candidate species for desert afforestation and dryland restoration. Its abundant flowering attracts a wide range of insects, supports honey production, and provides valuable forage resources for pollinators.
2. In this two-year study, we investigated the role of *Ziziphus spina-christi* in supporting parasitoid wasps across four sites that span a 200-600 mm annual rainfall gradient. Parasitoid abundance, diversity, community composition, and sugar-feeding frequency were assessed on *Z. spina-christi* and compared with surrounding vegetation.
3. Parasitoid abundance was consistently higher on *Z. spina-christi* than in the surrounding vegetation, especially during the hot and dry summer. Parasitoid diversity and community composition varied among sites along the aridity gradient. Mymarid wasps dominated some sites, mostly in autumn. Eulophids, braconids and scelionids were prominent in summer, and braconids became most dominant in autumn. The proportion of sugar-fed individuals increased during periods of greater flower availability.
4. These findings suggest that *Z. spina-christi* serves as an important nutritional resource and ecological refuge for parasitoid wasps, highlighting the potential of native drought-adapted plants to support beneficial insects and conservation biological control in dryland ecosystems.

## Introduction

Climate change is increasing the frequency and severity of droughts and heat waves worldwide, with particularly strong impacts in dryland ecosystems. Drylands cover approximately 40% of the Earth’s terrestrial surface and support a significant proportion of global biodiversity (Burrell et al., 2020). Given the current harsh abiotic conditions in drylands, further increases in aridity can alter species distributions, reduce resource availability, and disrupt ecological interactions. These changes may be especially challenging for populations of top predators, which are often regulated by bottom-up effects.

Parasitoid wasps, important top predators in insect food webs that depend on temporally variable resources, are abundant and diverse in drylands (Saabna & Keasar, 2024). They are among the most important natural enemies in terrestrial ecosystems, contributing significantly to the regulation of herbivorous insect populations and providing valuable biological control services (Fellowes et al., 2023). Despite their ecological importance, parasitoids face multiple challenges in arid environments, including limited host availability, scarcity of nutritional resources, and exposure to extreme temperatures and low humidity (Saabna & Keasar, 2024). As climate change intensifies environmental stress in dryland ecosystems, understanding the factors that promote parasitoid persistence becomes increasingly important. Characterizing dryland communities of parasitoids across space and time is a first step into understanding their adaptations to arid conditions.

Floral resources play a crucial role in the ecology of adult parasitoid wasps. Many species rely on nectar as a source of carbohydrates that support survival, dispersal, host-searching behavior, and reproduction (Fellowes et al., 2023). Consequently, the distribution and seasonal availability of flowering plants can strongly influence parasitoid populations and the ecosystem services they provide (Aartsma et al., 2020). In dryland environments, however, floral resources are often patchy, short-lived, and highly dependent on seasonal climatic conditions (Phillips et al., 2018), potentially limiting the persistence of parasitoid communities (Ratto et al., 2025).

Native perennial vegetation may help mitigate these limitations by providing reliable food resources and refuge habitats (Segoli et al., 2020). Such plants are increasingly recognized as important components of conservation biological control because they can sustain natural enemy populations when alternative resources are scarce. They are also less likely to become invasive than non-native plants. Drought-adapted plants may be particularly important in arid ecosystems, where they reliably provide ecological resources even under harsh environmental conditions.

One such species is *Ziziphus spina-christi* (L.) Desf., a widespread native tree in the Middle East that is highly tolerant of drought and heat stress. The species has long been recognized for its ecological, agricultural, and economic importance. The tree produces abundant flowers that attract a diverse insect community throughout the hot and dry summer and maintains vegetative cover throughout prolonged dry periods (Mizrahi & Nerd, 1996; Possingham, 1990; Vashishtha, 1997; Chovatia et al., 1993; Azam-Ali et al., 2006). Keasar and Shmida (2009) identified *Z. spina-christi* as an important forage plant that produces nectar and strongly attracts honey bees, small bees and flies, particularly when alternative floral resources are limited. It supports honey production and provides valuable forage resources for pollinators and livestock. These traits make *Z. spina-christi* a promising candidate species for desert afforestation, dryland restoration, and sustainable land management programs aimed at combating land degradation and desertification.

*Z. spina-christi’*s ability to provide nectar, shelter, and favorable microhabitats suggests that it may serve as an important resource for parasitoid wasps in water-limited environments. Furthermore, by supporting populations of natural enemies, *Z. spina-christi* may contribute to conservation biological control in surrounding agricultural landscapes. Yet, despite the potential ecological importance of *Z. spina-christi* and other native drought-adapted trees, their role in supporting parasitoid communities across gradients of aridity remains poorly understood. In particular, little is known about how the trees’ floral resources influence parasitoid abundance, diversity, community composition, and nutritional ecology in dryland ecosystems.

In this study, we investigated the role of *Ziziphus spina-christi* in supporting parasitoid wasp communities across sites that differ in aridity. Specifically, we examined parasitoid abundance, diversity, species composition, and sugar-feeding associated with *Z. spina-christi* to evaluate its potential importance as a nutritional resource and ecological refuge for parasitoids in dryland environments.

## Methods

### Parasitoid sampling

We characterized the parasitoid community on *Z. spina-christi* trees across sites and seasons over two years (October 2023 - August 2025). To compare sites, we sampled natural populations of trees in four locations in northern Israel (Fig. 1) along a rainfall gradient in July, when the trees were in peak bloom: Tirat Tzvi (200–300 mm annual rainfall), Reshafim (300–400 mm), Nahal Tavor (400–500 mm), and Khawaled (500–600 mm). Additional sampling was conducted during autumn (October), winter (January), and spring (April) in Nahal Tavor and Reshafim to compare parasitoid assemblages between seasons. Insects were collected from fifteen *Z. spina-christi* trees in each site using a Vortis insect suction sampler. Each tree was sampled for 60 seconds to standardize sampling effort. Two 30-second samples were collected from every tree: one sample was collected into dry vials, kept on ice, and later used to evaluate the parasitoids’ sugar-feeding frequency (see section ‘Anthrone tests’ below). The second sample was preserved in 70% ethanol for parasitoid identification and abundance analyses. All samples were transported on ice and stored at −20°C until processing. Flowering density on the day of sampling was estimated by counting the number of inflorescences on a 2-m branch at breast height in each tree. Flowers were counted in ten randomly selected inflorescences, and the mean number of flowers per inflorescence was multiplied by the total number of inflorescences on the branch to estimate flower abundance. Additionally, in 2025 only, the herbaceous vegetation surrounding each sampled tree was suction-sampled for 15 seconds. The surrounding vegetation was sampled in all four field sites in summer, during the trees’ peak flowering. To detect possible seasonal trends in the use of *Ziziphus* trees vs. herbaceous vegetation, we also suction-sampled the vegetation in spring in two of the study sites (Reshafim and Nahal Tavor). To standardize sampling effort and enable direct comparisons with tree samples, parasitoid counts from the surrounding vegetation were multiplied by two, yielding values equivalent to a 30-second sampling period.

**Figure 1.**
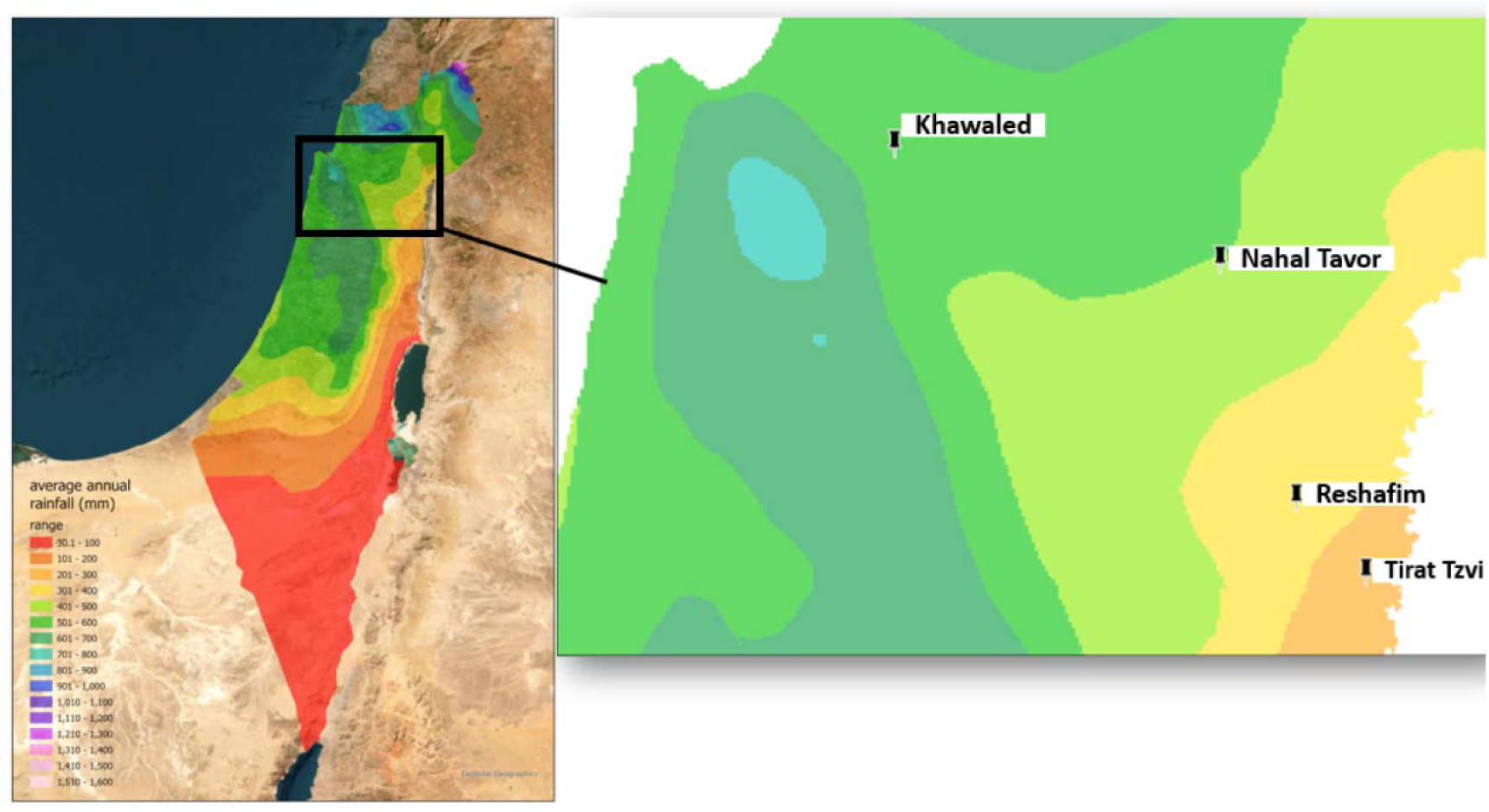
Locations of the four study sites along a mean annual rainfall gradient in northern Israel: Tirat Tzvi (200–300 mm), Reshafim (300–400 mm), Nahal Tavor (400–500 mm), and Khawaled (500–600 mm). Colors represent mean annual rainfall (mm); black markers indicate the study sites. The inset map shows the location of the study region within Israel.

### Sorting of samples

The abundance of all parasitoids in the samples was determined. Parasitoids were identified at the family level. Dominant parasitoids (defined as species that made up >3% of all individuals) were further identified to species.

### Anthrone tests

Sugar-feeding was assessed in the parasitoids from the dry samples using qualitative cold anthrone assays, which detect fructose and sucrose in the parasitoid gut (Heimpel et al., 2004). Individuals showing a color response within one hour were recorded as anthrone positive, indicating recent sugar feeding. One researcher assayed, scored as anthrone-positive or negative, and photographed all individuals. A second researcher validated the scores based on the photo images.

#### Data analysis

##### Parasitoid abundance and community composition

Parasitoid abundance was analyzed using a generalized linear mixed model (GLMM), with site, year, season, and habitat as fixed effects and tree identity included as a random effect to account for repeated sampling of the same trees. The poisson family with a log link function, which is suitable for count data, was used in the model.

Shannon diversity indices were calculated from parasitoid family abundances for each sample. The effects of site, year, and season on family-level diversity were tested using a linear mixed-effects model with a gaussian data distribution and an identity link function, with tree included as a random effect.

Community composition of parasitoid families was analyzed using non-metric multidimensional scaling (NMDS) based on Bray–Curtis dissimilarity matrices calculated from family-level abundance data. Ordinations were used to visualize variation in parasitoid assemblages among sites and seasons. Environmental fitting was applied to identify the parasitoid families most strongly associated with observed compositional gradients. We used permANOVA tests to determine the effects of site and season on the parasitoids’ family-level composition.

##### Anthrone tests

The proportion of anthrone-positive individuals per tree was used to estimate sugar consumption frequencies across seasons, years, and study sites. We used a Beta regression, suitable for proportions data, to evaluate the effects of season, year, and site on sugar feeding frequencies.

We used R version 4.6.0 (R Core Team, 2026) for data visualization and analyses. We used packages lme4 (Bates et al., 2015) for the Generalized Linear Models, vegan (Oksanen et al., 2026) for the NMDS and permANOVA analyses, and betareg (Cribari-Neto & Zeileis, 2010) for the beta regression.

## Results

### Parasitoid abundance

Parasitoid abundances differed significantly among sites (GLMM: χ²₃ = 16.23, P = 0.001), years (χ²₂ = 644.05, P < 0.001), seasons (χ²₃ = 666.44, P < 0.001), and habitats (χ²₁ = 119.90, P < 0.001) (Figs. 2-4). After accounting for site, year, season, and repeated sampling of individual trees, the estimated parasitoid abundance was significantly higher on *Ziziphu*s trees (estimated mean = 4.30, 95% CI: 2.95–6.29) than in the surrounding vegetation (estimated mean = 1.65, 95% CI: 1.09–2.48) (Fig. 2). Parasitoid abundance in the surrounding vegetation was 0.382 times that on *Ziziphus* trees (z = −10.95, P < 0.0001), corresponding to approximately 2.6-fold greater abundance on *Ziziphus*. This habitat effect was significantly weaker in spring than in summer (Season × Habitat: β = −1.356, SE = 0.188, z = −7.227, P < 0.001, Fig. 3). Khawaled, the most mesic site, had the lowest abundance of parasitoids. Relative to Khawaled, abundance was significantly higher at Nahal Tavor (β = 0.713, P = 0.003), Reshafim (β = 0.658, P = 0.006), and Tirat Tzvi (β = 0.929, P < 0.001) (Fig. 2). Parasitoid abundance was lowest in winter and gradually increased in spring and summer (Fig. 4).

**Figure 2.**
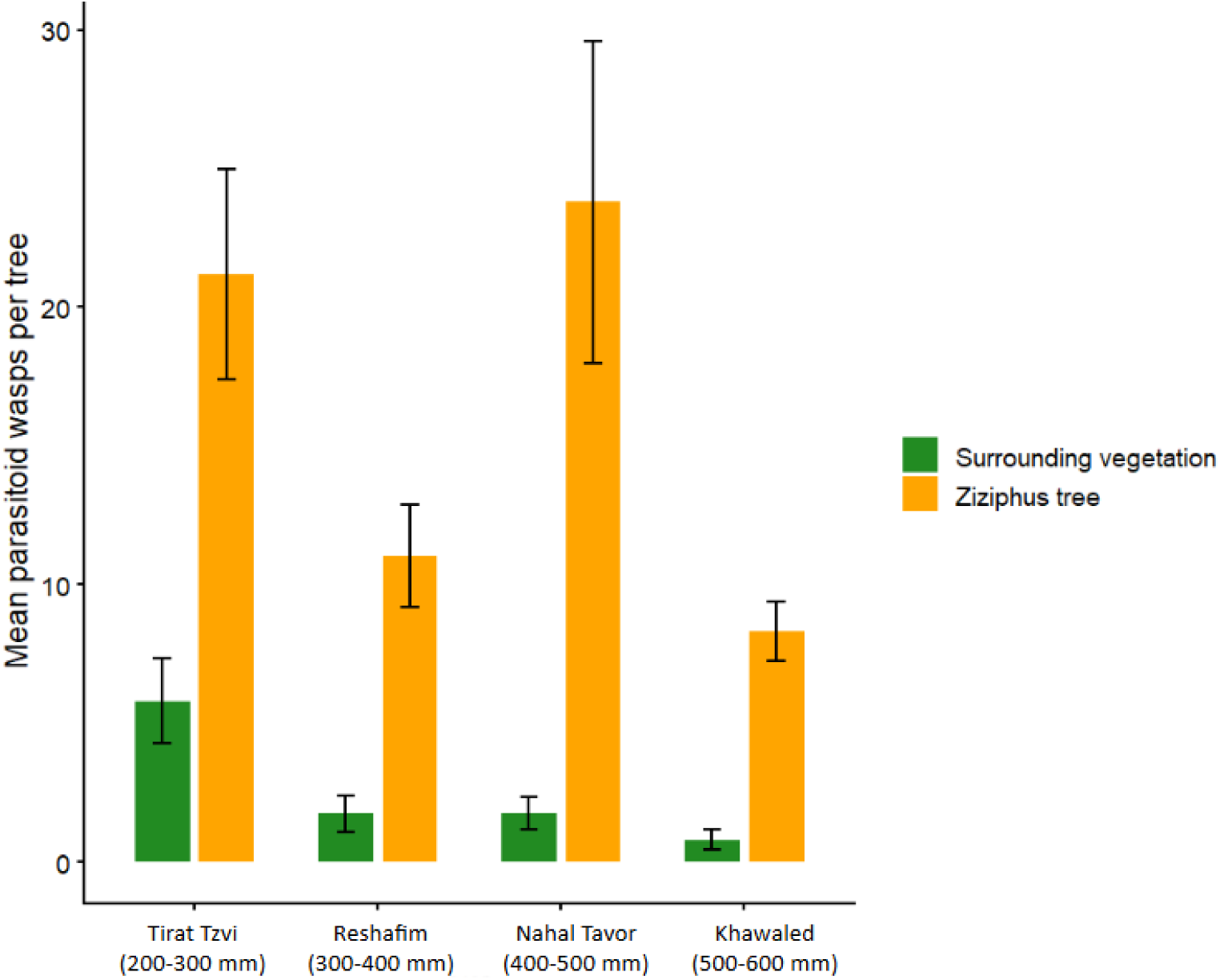
Mean abundance (± SE) of parasitoid wasps collected from *Ziziphus spina-christi* trees in 2024-2025 and surrounding vegetation (in 2025 only) across four study sites.

**Figure 3.**
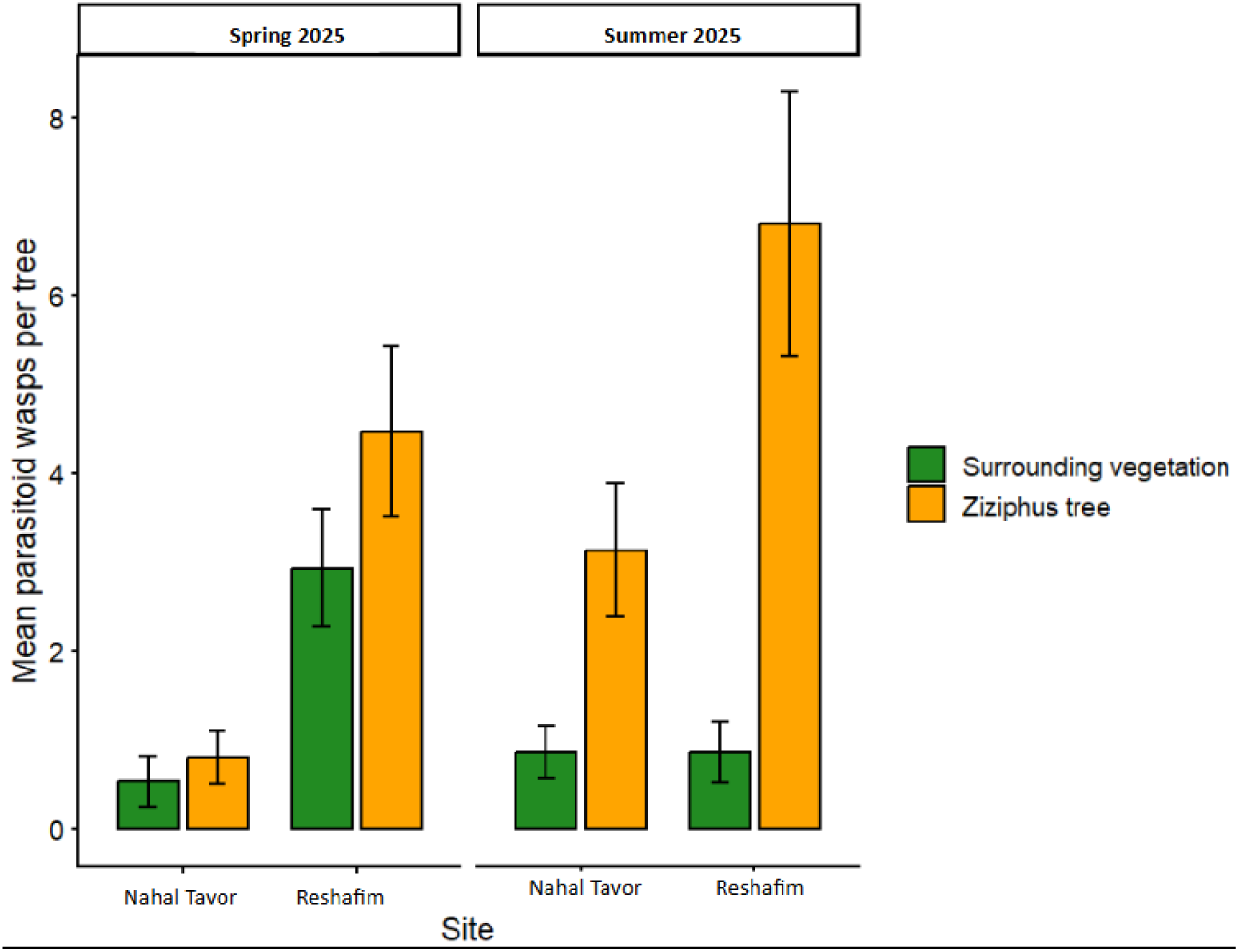
Mean number of parasitoid wasps collected per tree from *Ziziphus spina-christi* trees and the surrounding vegetation at Nahal Tavor and Reshafim during spring and summer 2025.

**Figure 4.**
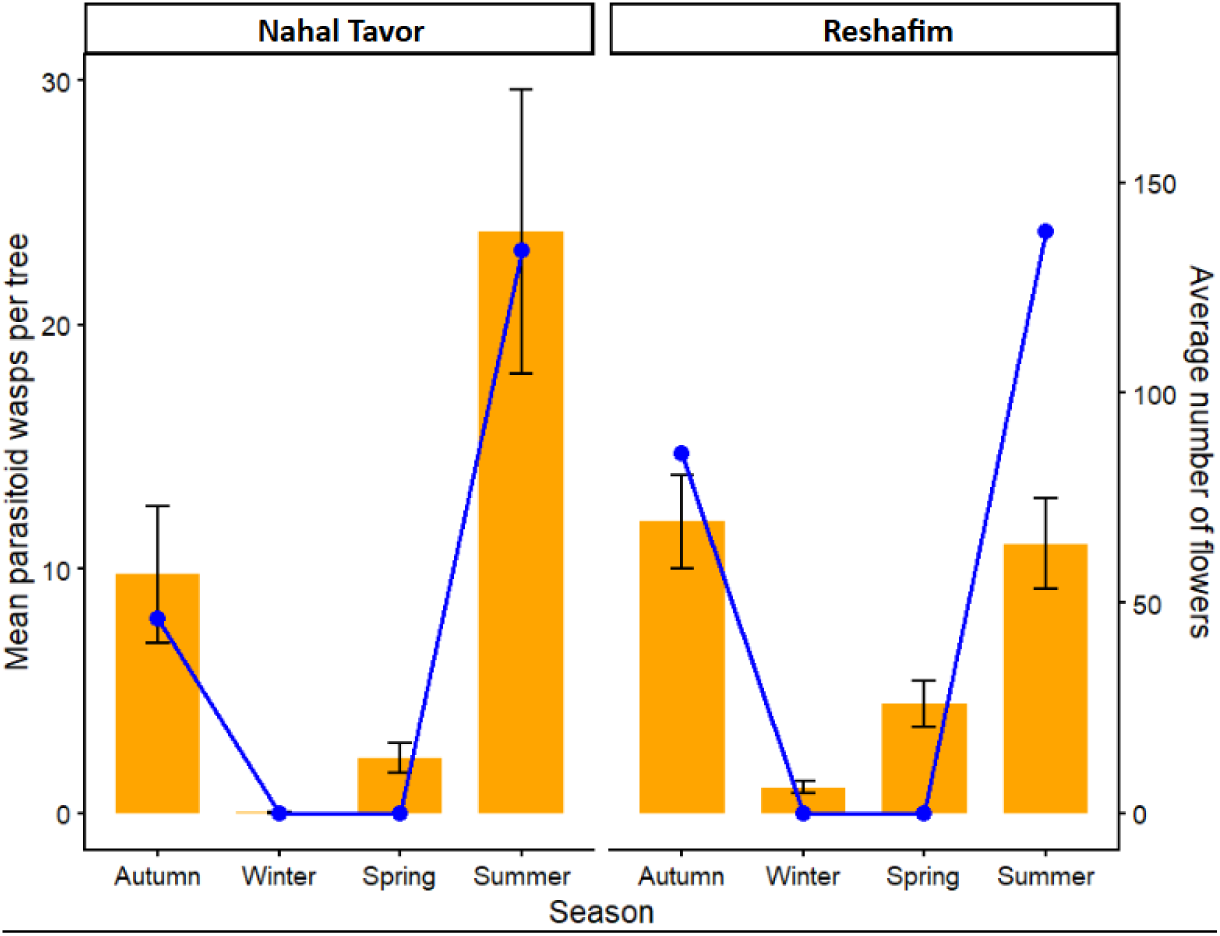
Average abundance of parasitoid wasps collected from trees (bars; mean ± SE) and average number of flowers (line) at Nahal Tavor and Reshafim across different seasons in 2024 and 2025.

### Anthrone-positive parasitoid patterns

The proportion of anthrone-positive parasitoids varied seasonally (Fig. 5). Beta regression indicated that the proportion of sugar-fed parasitoids was significantly lower in spring than in autumn (β = −1.247, z = −2.710, P = 0.0067), whereas summer (P = 0.578) and winter (P = 0.821) did not differ significantly from autumn. The trees were not in bloom in winter and spring. Nevertheless, a considerable proportion of the parasitoids collected from the trees in the winter and spring scored positive for anthrone (Fig. 5). No significant difference was detected between Nahal Tavor and Reshafim (P = 0.561), and year effects were also not significant (P=0.1587).

**Figure 5.**
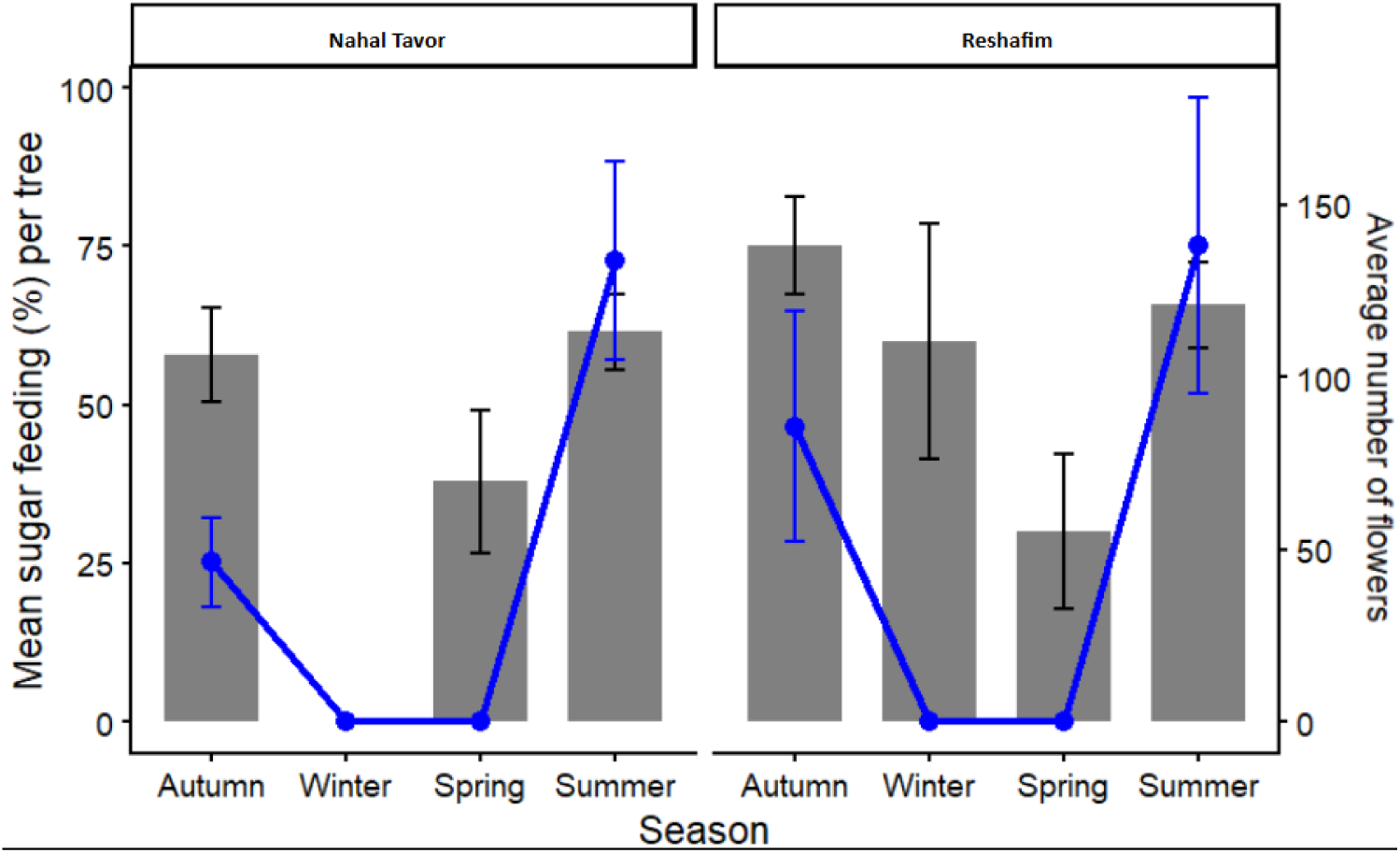
Proportion of anthrone-positive parasitoids (bars; mean ± SE) and average flower number per tree (line) across seasons in 2024–2025.

#### Parasitoid diversity

##### Family-level parasitoid diversity among sites

During summer, parasitoid family diversity differed significantly among the four study sites (linear mixed-effects model: χ²₃ = 10.98, P = 0.012), after accounting for year and repeated sampling of individual trees. The estimated Shannon diversity was highest at Khawaled (1.00, 95% CI: 0.80–1.21) and Reshafim (0.97, 95% CI: 0.76–1.18), followed by Tirat Tzvi (0.72, 95% CI: 0.51–0.93), and was lowest at Nahal Tavor (0.58, 95% CI: 0.37–0.79). Shannon diversity at Nahal Tavor was significantly lower than at Khawaled (P = 0.025) and Reshafim (P = 0.047). No other pairwise differences among sites were significant (Tukey-adjusted comparisons, all P ≥ 0.230). Diversity did not differ between 2024 and 2025 (χ²₁ = 0.03, P = 0.864, Fig. 6)

**Figure 6.**
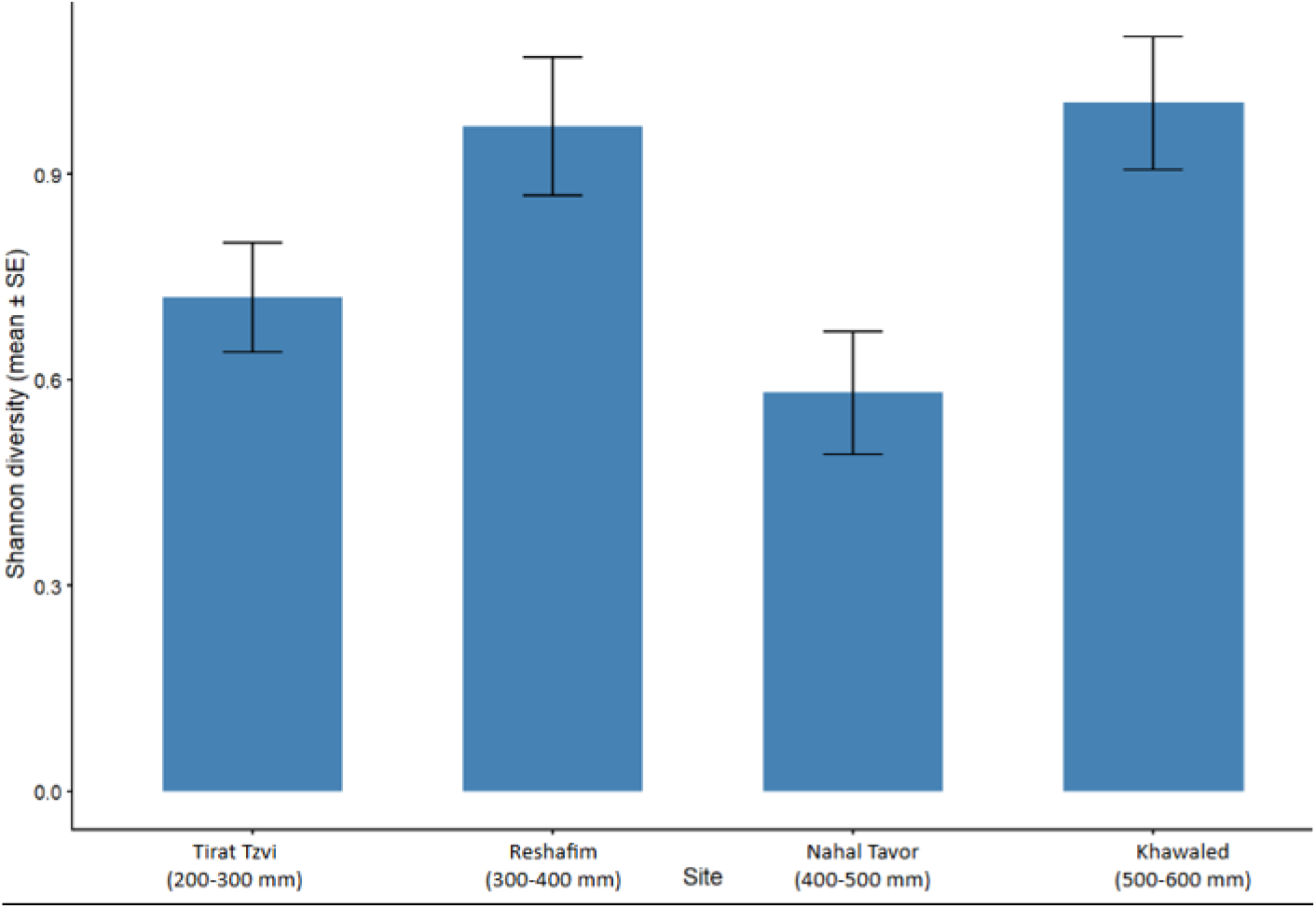
Shannon diversity index (mean ± SE) of parasitoid wasp communities at the family level across four sites along a rainfall cline in 2024–2025.

##### Seasonal variation in parasitoid diversity

Reshafim and Nahal Tavor were sampled in different seasons. In these sites, Shannon diversity differed significantly among seasons (linear mixed-effects model: χ²₃ = 29.19, P < 0.001, Fig. 7) as well as between sites (χ²₁ = 7.44, P = 0.006). The site-by-season interaction was not significant (χ²₃ = 2.79, P = 0.426), indicating that the overall seasonal pattern was similar at both sites. The effect of year was also not significant (χ²₂ = 4.99, P = 0.082).

At both Reshafim and Nahal Tavor, diversity was lowest during winter. At Nahal Tavor, winter diversity was significantly lower than in autumn (P = 0.0006), spring (P = 0.010), and summer (P < 0.0001). Similarly, at Reshafim, winter diversity was significantly lower than in autumn (P < 0.0001), spring (P = 0.0002), and summer (P < 0.0001). No significant differences were detected among autumn, spring, and summer at either site.

Estimated Shannon diversity was higher at Reshafim than at Nahal Tavor during autumn (1.18 vs. 0.71; P = 0.006), spring (1.02 vs. 0.50; P = 0.001), and summer (1.10 vs. 0.71; P = 0.0025). The difference between the two sites was not significant during winter (0.34 vs. 0.13; P = 0.200, Fig. 7).

**Figure 7.**
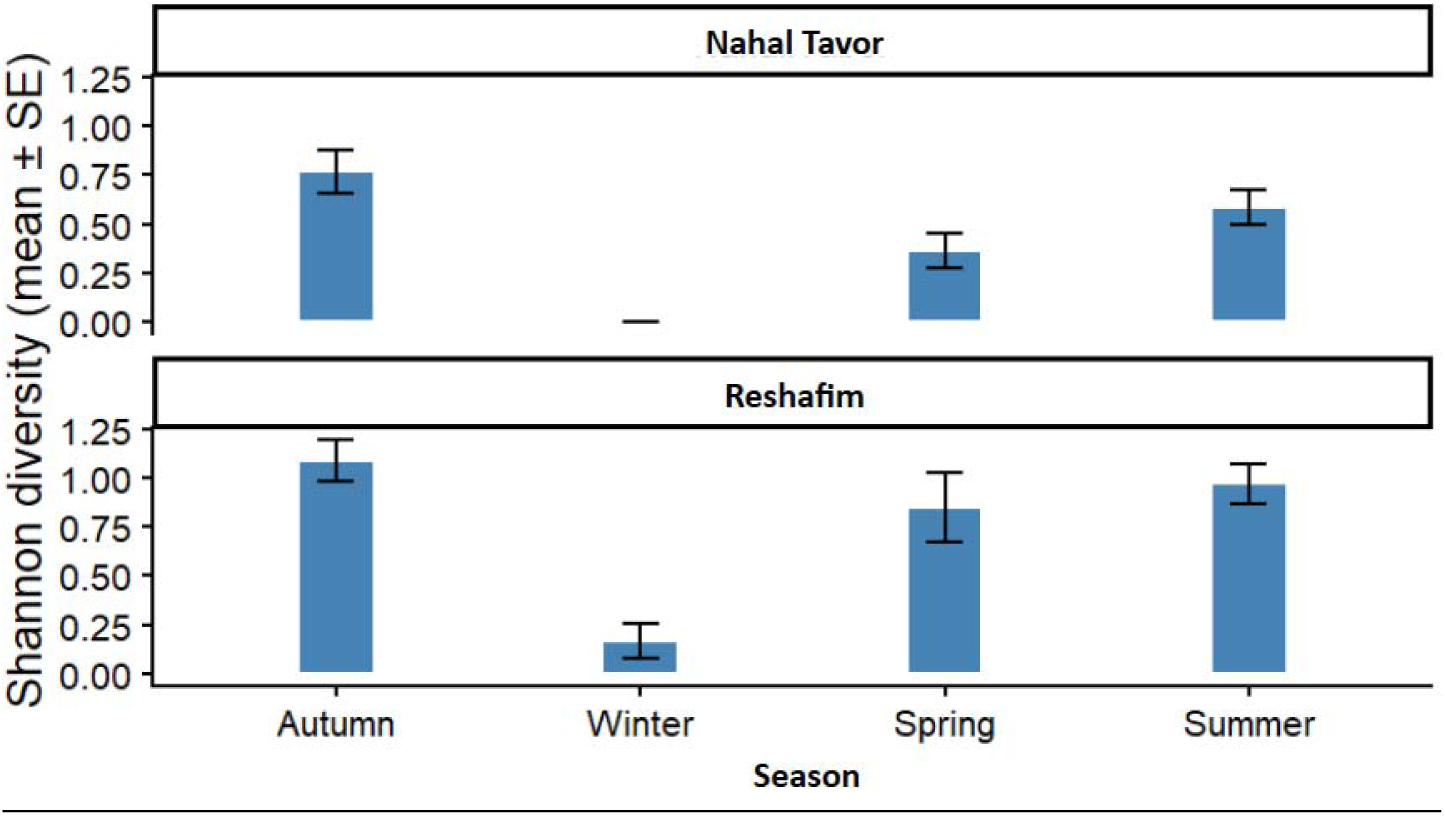
Shannon diversity index (mean ± SE) of parasitoid wasp communities at the family level across seasons at Nahal Tavor and Reshafim in 2024–2025.

##### Family-level community composition

NMDS ordination based on Bray–Curtis dissimilarities showed variation in parasitoid family composition among the four study sites (Fig. 8). Although samples overlapped considerably, some separation was evident, and PERMANOVA confirmed significant differences among sites (pseudo-F₃,₈₀ = 5.13, R² = 0.161, p = 0.001; 999 permutations). Site explained 16.1% of the variation in family composition. The site ordination suggested that samples from Khawaled were more strongly associated with Mymaridae and Braconidae than samples from the remaining sites, although families were mixed across sites.

Seasonal variation was also significant (pseudo-F₂,₆₅ = 8.30, R² = 0.203, p = 0.001; 999 permutations), with season explaining 20.3% of the variation. In the seasonal ordination, autumn samples were associated with Mymaridae and Braconidae, summer samples with Eulophidae, Scelionidae, and Braconidae, while spring samples comprised diverse parasitoid families.

**Figure 8.**
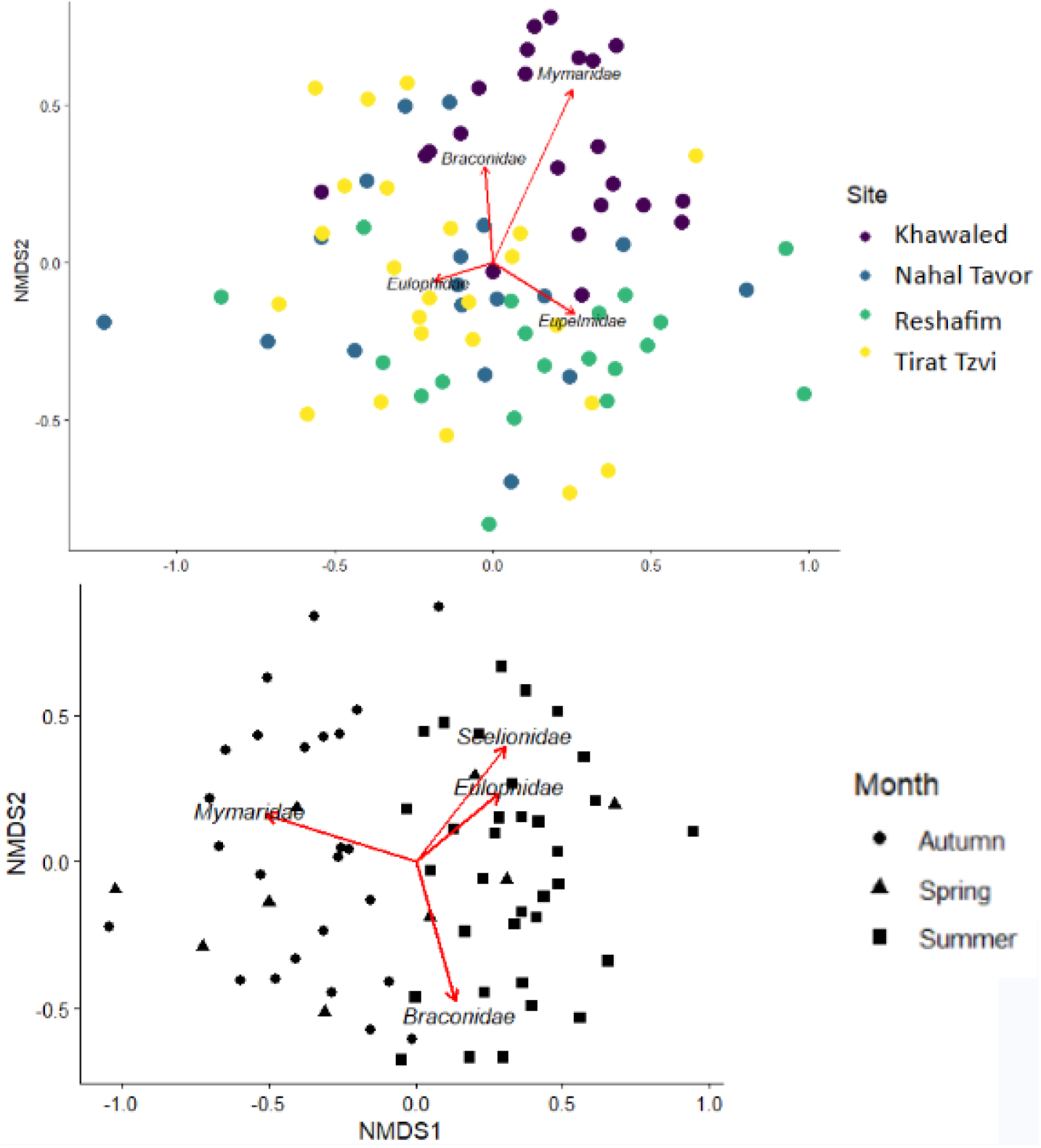
NMDS ordinations of parasitoid family composition based on Bray–Curtis dissimilarities. Top: family composition by site. Bottom: family composition by season. Only trees with ≥ 5 collected parasitoids were included in the analyses. Arrows represent family vectors showing the taxa most strongly associated with patterns of community variation.

##### Dominant parasitoids

The parasitoid assemblage was dominated by the eulophid *Zagrammosoma ramotensis* and by the braconids, *Gnamptodon avigail* and *Glyptapanteles sp*. The relative composition of these dominant parasitoids differed substantially among the four study sites (Fig. 9). *Z. ramotensis* was particularly dominant at Nahal Tavor, where it represented approximately 66.9% of the parasitoid assemblage, and at Tirat Tzvi, where it accounted for approximately 57.5%. In Khawaled and Reshafim, *Z. ramotensis* represented only 12.8% and 28% of all parasitoids, respectively.

*G. avigail* also showed pronounced variation among sites. It accounted for 18.4% of parasitoids at Reshafim and 10.2% at Tirat Tzvi, but represented only 4.1% at Khawaled and was not represented among the dominant taxa at Nahal Tavor. *Glyptapanteles sp.* constituted a relatively small proportion of the assemblage across all sites. Seasonally, the dominant species were observed mainly in summer and were largely absent during the other seasons.

Overall, these patterns indicate marked spatial variation in the identity and relative dominance of parasitoid taxa across the rainfall gradient (Fig. 9).

**Figure 9.**
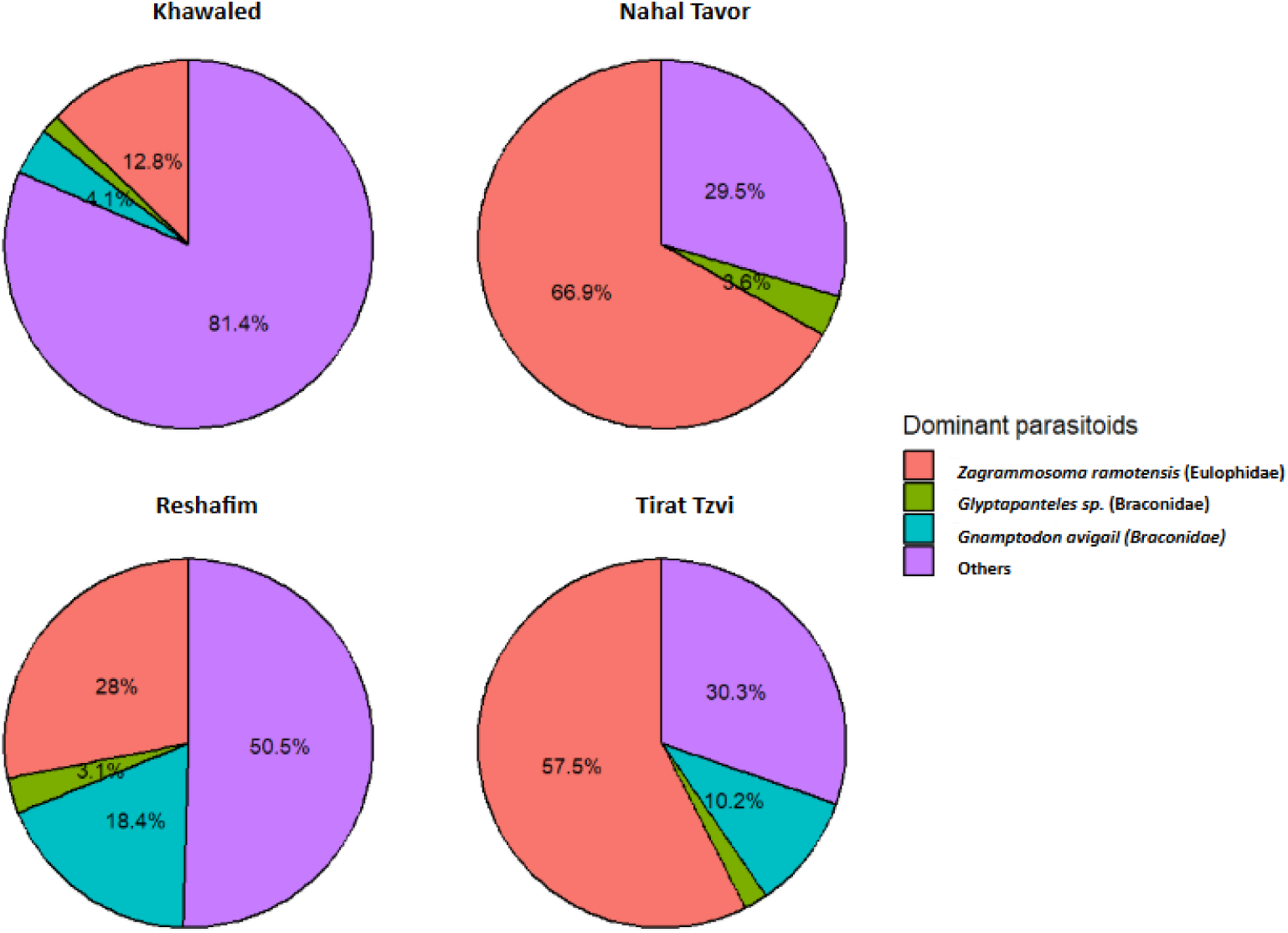
Mean abundance (± SE) of dominant parasitoid species collected across the four study sites during 2024–2025

## Discussion

This study characterized communities of parasitoid wasps on *Z. spina-christi* trees along a rainfall gradient. We asked whether parasitoids are more abundant on the trees than in other nearby habitats. We found, indeed, higher parasitoid abundances on *Ziziphus spina-christi* trees than in the surrounding vegetation across all four study sites (Fig. 2-3). This pattern suggests that *Ziziphus* trees provide important resources for parasitoids, potentially including nectar, shelter, and hosts. These resources may be particularly valuable in dry environments, where alternative food sources and vegetation can be limited (Phillips et al., 2018). The difference in parasitoid abundance between *Ziziphus* trees and surrounding vegetation increased in summer, suggesting that the trees may be especially important to parasitoids during this season. While the trees reached peak flowering in summer (Fig. 5), the surrounding herbaceous plants typically flower in spring and dry out during the summer. This seasonal pattern raises the question of whether flowering, in particular, contributes to parasitoid activity and feeding.

Our sugar-feeding results provide some insight into this possibility. The proportion of anthrone-positive parasitoids did not differ significantly between winter, summer, and autumn, although the trees did not flower in winter (Fig. 5). This suggests that parasitoids may also use other sugar sources, such as honeydew, and that feeding may be influenced by additional factors.

Seasonal changes were evident in parasitoid abundance, which was relatively high in summer and autumn but very low in winter (Fig. 4). At Nahal Tavor, parasitoid abundance peaked during peak *Ziziphus* flowering. At Reshafim, however, abundance did not show the same summer peak, possibly because harsh summer conditions continued to limit parasitoid activity despite the trees’ flowering. This contrast strengthens the evidence that flowering may support parasitoid activity, but that its effect depends on local environmental conditions, including temperature and host availability (Aartsma et al., 2020).

Comparisons across seasons and sites indicate the parasitoid diversity did not simply mirror abundance. Seasonally, family-level diversity was lowest in winter and generally higher in summer and autumn at both Nahal Tavor and Reshafim (Fig. 7). However, summer abundance was much higher than autumn abundance in Nahal Tavor, without a corresponding difference in diversity (Fig. 4). Thus, favorable seasonal conditions may increase the numbers of some parasitoid families without increasing overall community diversity to the same extent.

Spatially, parasitoid abundance and diversity peaked in different sites along the rainfall gradient. Nahal Tavor and Tirat Tzvi showed higher abundance than Reshafim and Khawaled (Fig. 2), whereas Reshafim and Khawaled had higher family-level diversity (Fig. 6). This difference between abundance and diversity reinforces the between-season comparison, indicating that a large number of parasitoids does not necessarily indicate a more diverse community. Neither abundance nor diversity varied monotonously with aridity across sites. This suggests that rainfall alone does not explain the observed patterns; local vegetation, host availability, and other environmental conditions may also influence parasitoid communities.

Differences among sites were also reflected in family-level community composition. PERMANOVA tests showed significant differences among sites and seasons, although samples overlapped considerably in the NMDS ordination (Fig. 8). Site explained 16.1% of the variation, while season explained 20.3%, indicating that seasonal differences accounted for a somewhat larger proportion of variation in this analysis.

The family-level patterns were also reflected in the dominant parasitoid taxa (Fig. 9). *Zagrammosoma ramotensis* (Eulophidae) was particularly dominant at Nahal Tavor (66.9% of all parasitoids) and Tirat Tzvi (57.5%). This species is a larval ectoparasitoid associated with leaf-mining Lepidoptera and Diptera and is considered an idiobiont, meaning that host development stops following parasitism (Yefremova & Kishinevsky., 2021) (Cao et al., 2014). This pattern is consistent with the NMDS ordination, which showed an association between Eulophidae and samples from Nahal Tavor. *Gnamptodon avigail* (Braconidae) was more prominent at Reshafim (18.4%) and Tirat Tzvi (10.2%). This is a recently described species, and little is currently known about its ecology and host associations (Pisanty et al., 2026). However, members of the genus *Gnamptodon* are generally associated with leaf-mining Lepidoptera and are considered koinobiont larval endoparasitoids, allowing host development to continue after parasitism. (Pisanty et al., 2026). *Glyptapanteles sp.* (Braconidae) was also recorded. Members of this genus are typically koinobiont larval endoparasitoids of Lepidoptera, developing inside caterpillar hosts while allowing them to continue growing (Arias-Penna et al., 2019). At Khawaled, most parasitoids belonged to other taxa (81.4%). This high proportion of “other” taxa is consistent with the higher family-level diversity observed at Khawaled, suggesting that its community included a wider range of less dominant taxa. These site-specific patterns may reflect differences in host availability, vegetation, or local environmental conditions.

Taken together, the results suggest that *Ziziphus* spina-christi may serve as an important resource and refuge for parasitoid communities in dryland environments, particularly during summer, when the abundance difference between the trees and surrounding vegetation was greatest. At the same time, parasitoid feeding, abundance, diversity, and community composition varied across seasons and sites, indicating that the role of *Ziziphus* is shaped by more than flowering alone. Maintaining native *Ziziphus* trees in dryland agricultural landscapes may support conservation biological control, but further research is needed to determine whether parasitoids associated with these trees improve pest suppression in nearby crops.

## Supporting information

Link to data and R code

## Acknowledgements

The study was funded in part by the Middle East Regional Cooperation (MERC) Program, grant No. SIS70023GR0011 from the United States Department of State and the United States Agency for International Development. Additional funding was provided by the Israel Council of Higher Education, grant ‘Biodiversity monitoring in Israel’s agroecosystems’. We used the AI tool ChatGPT to improve the flow and clarity of the text. We thank Ronen Shapir, Soliman Saabna and Tom Siyach for their help with field sampling and sorting of the samples.

